# Prenatal Carotenoid Supplementation with Lutein or Zeaxanthin Ameliorates Oxygen-Induced Retinopathy (OIR) in *Bco2^-/-^* Macular Pigment Mice

**DOI:** 10.1101/2022.12.28.521962

**Authors:** Ranganathan Arunkumar, Binxing Li, Emmanuel K. Addo, M. Elizabeth Hartnett, Paul S. Bernstein

## Abstract

**Purpose:** Premature infants at risk of retinopathy of prematurity (ROP) miss placental transfer of carotenoids, lutein (L) and zeaxanthin (Z), during the third trimester. We previously demonstrated that prenatal L and Z supplementation raises carotenoid levels in infants at birth in the <u>L</u>utein and <u>Z</u>eaxanthin in <u>P</u>regnancy (L-ZIP) study (NCT03750968). Based on their antioxidant effects and bioavailability, we hypothesized that prenatal maternal supplementation with macular carotenoids would reduce risk of ROP. To test this hypothesis, we utilized “macular pigment mice” genetically engineered to take up L and Z into the retina in a model of oxygen-induced retinopathy (OIR).

**Methods:** Pregnant *Bco2*^*-/-*^ mice were divided into nine experimental subgroups based on the type of supplementation (L, Z, or placebo) and on maternal supplementation start date corresponding to the three trimesters of human fetal development (E0, E11, and P1). Pups and nursing mothers were exposed to 75% O_2_ for 5 days (P7-12) and returned to room air for 5 days (P12-17). Pups were sacrificed at P12 and P17, and their retinas were analyzed for vaso-obliteration (VO) and intravitreal neovascularization (INV).

**Results:** Pups of pregnant mice supplemented with L or Z had significant reductions in VO and INV areas compared to placebo. Prenatal carotenoid supplementation starting at E0 or E11 was significantly more protective against OIR than postnatal supplementation starting at P1.

**Conclusion:** Prenatal supplementation with L and Z was beneficial in a mouse OIR model. We recommend testing prenatal L and Z supplementation in future human clinical trials to prevent ROP.

## Introduction

Retinopathy of prematurity (ROP) is a retinal vasoproliferative disease that affects premature babies that is one of the most common causes of childhood blindness. ^1^ The pathogenesis of ROP is multifactorial, and its major risk factors are low birth weight, young gestational age, oxygen stresses, and nutrition. ROP has been described by a two-phase hypothesis based on animal models of oxygen-induced retinopathy (OIR). The first phase is characterized by disruption of normal retinal vascular growth caused by fluctuations in oxygenation, hyperoxia-induced damage to newly developed capillaries, oxidative stress, and other preterm complications. This is followed by hypoxia-induced neovascularization that may lead to retinal detachment and vision loss. ^2,3^ In a normal human fetus, retinal development starts from the optic nerve and proceeds toward the periphery at ∼16 weeks of gestation and is completed around 36 weeks of gestation. ^4^ Preterm infants with an immature cardiopulmonary system may require oxygen therapy, but oxygen stresses can lead to areas of avascular retina, which can become hypoxic as the infants are transitioned to room air. This is believed to lead to hypoxia-induced pathological vasoproliferation. ^5^

Oxidative stress is a major risk factor in the development and progression of ROP ^6^ because life-saving therapies in neonatal intensive care units (ICUs) may expose premature infants to high oxygenation, intense light, multiple medications, and stressful conditions. Also preterm babies are more susceptible to oxidative stress because the protective antioxidant system of the immature retina has insufficient ability to quench reactive oxygen species. ^7^ Lutein (L) and zeaxanthin (Z) are natural antioxidants commonly present in human diets from fruits, vegetables, and egg yolks. L, Z, and their ocular metabolite *meso*-zeaxanthin (MZ), are collectively known as the macular pigment (MP) and are preferentially localized in the foveal region of the human macula, an area specialized for central and distinct spatial vision. MP protects the macula from light-induced oxidative damage through their blue light filtering properties and their antioxidant and anti-inflammatory activities. ^8, 9^

L and Z may play a crucial role in prenatal life, as evidenced by their presence in the placenta and cord blood at the time of birth due to placental transfer of carotenoids from the mother to the fetus especially during the third trimester. ^10^ The presence of macular carotenoids in prenatal eyes as early as 17-22 weeks of gestation suggests a potential role in the development of the primate fovea and other important retinal structures. ^11, 12, 13^ The MP carotenoids continue to accumulate in the human retina from birth until at least 7 years of age. ^14^ Among the various dietary carotenoids, L and Z are the most abundant ones in breast milk, and their deficiency may negatively influence an infant’s health. ^15^ Furthermore, it has been observed that the mother’s carotenoid status may be depleted during pregnancy as she transfers these nutrients to her child. ^16^ Premature babies miss the third-trimester placental transfer of carotenoids, and they may rapidly become further carotenoid depleted due to their exposure to severe oxidative stress. ^10^ Moreover, currently available prenatal vitamins and premature infant nutritional formulations typically do not have any added L or Z, ^17^ but we have recently shown that L and Z in human prenatal supplements started before 14 weeks of gestation at an AREDS2 dose of 10 mg/day and 2 mg/day, respectively, safely increased the systemic and ocular carotenoid status of the mother and her newborn infant (The <u>L</u>utein and <u>Z</u>eaxanthin in <u>P</u>regnancy (L-ZIP) study; NCT03750968). ^18^

Published clinical studies of the potential benefit of L and Z supplements for the prevention of ROP have focused on postnatal infant supplementation to preemies at concentrations comparable to an AREDS2 dose on a mg/kg basis. These studies did not show a statistically significant decrease in disease progression, ^19, 20, 21,^ but we speculate that these interventions may have been administered too late. Therefore, we hypothesize that prenatal maternal supplementation with macular carotenoids given to the mother at an AREDS2 dose could alleviate ROP pathology better than prior postnatal studies.

Before proceeding to human clinical trials of prenatal L and Z for prevention of ROP, it is important to perform efficacy tests in a suitable animal model that actively accumulates dietary carotenoids into the retina. The mouse OIR model is a well-established, consistent, and economical model to mimic aspects of the two-phased hypothesis: Phase I, in which there is damage to newly developed capillaries, sometimes termed vaso-obliteration (VO); and Phase II, in which there is intravitreal neovascularization (INV). Retinal vascular development occurs *ex utero* in mice because they are born with incomplete retina vasculature at full-term birth. Until recently, however, the potential efficacy of carotenoids against mouse OIR was not testable because wild-type (WT) mice do not accumulate retinal carotenoids even in the face of high-dose supplementation. To address this problem of poor retinal uptake of carotenoids, we previously developed transgenic “macular pigment mice” whose xanthophyll cleavage enzyme, beta-carotene oxygenase 2 (Bco2), has been knocked out. *Bco2*^*-/-*^ mice readily accumulate L and Z in their retinas in response to oral supplementation. ^22, 23^ Here, we use these macular pigment mice to test the relative efficacies of prenatal and postnatal L and Z in a mouse model of OIR.

## Materials and Methods

### Materials

L and Z HPLC standards (≥ 98% pure) were provided by Kemin Health (Des Moines, IA, USA) and DSM Nutritional Products, Ltd. (Kaiseraugst, Switzerland), respectively. L, Z, and placebo Actilease® beadlets were supplied by DSM Nutritional Products, Ltd. (Kaiseraugst, Switzerland). L and Z Actilease® beadlets were mixed with a previously reported base diet (Testdiet®, Richmond, IN, USA) at a dosage of 1 g/kg (∼2.6 mg per mouse per day) ^24^. All organic solvents were of HPLC grade and were purchased from Thermo Fisher (Waltham, MA, USA).

### Animals

*Bco2*^*-/-*^ breeding pairs on a C57BL*/6/J* background were received from Case Western Reserve University, Cleveland, OH. *Bco2*^*-/-*^ mice were provided food and water *ad libitum* and kept on a 12-hour light/dark cycle. Animal experiments were designed and performed according to the guidelines and regulations of the ARVO Statement for the Use of Animals in Ophthalmic and Vision Research. Experimental animal protocols were approved by the Institutional Animal Care and Use Committee of the University of Utah.

Pregnant mice were supplemented L and Z through Actilease® beadlets in a base diet as described above providing *in utero* supplementation to fetuses. Pups were nourished postnatally through maternal milk.

### Oxygen-induced retinopathy (OIR)

Female *Bco2*^*-/-*^ mice were mated with male *Bco2*^*-/-*^ mice, and pregnancy was confirmed based on the presence of a vaginal plug at embryonic day 0 (E0). Pregnant *Bco2*^*-/-*^ mice were divided into three experimental groups based on the start date of supplementation (E0, E11, and postnatal day 1 (P1), dates chosen to be comparable to the beginnings of the three trimesters of human pregnancy). Each experimental group was further divided into three supplementation subgroups (L, Z, or placebo diets). At every evaluation timepoint (P12 and P17), each subgroup had at least 3 pups from at least two different litters. Oxygen-induced retinopathy (OIR) was induced in newborn mice as previously described. ^25^ Briefly, on postnatal day 7 (P7) when the inner vascular plexus covers the retinal extent, *Bco2*^*-/-*^ mouse litters from all three groups along with their nursing mothers were exposed to 75% oxygen for 5 days (until P12) in a ProOx model P360 hyperoxia chamber (Biosphere Ltd., Parish, NY, USA) to initiate OIR and then returned to room air (21% oxygen) for another 5 days (until P17) for vascular regrowth and intravitreal neovascularization (INV) until pup sacrifice at P12 or P17. A schematic summary of the research protocol is provided in **Supplementary Table 1**. Although the number of pups per litter tends to be smaller whenever carotenoid supplementation is started prenatally in *Bco2*^*-/-*^ mice, **Supplementary Figure 1** shows that pups in our various subgroups had identical weight gain at P7, P12, and P17.

Mouse pups were sacrificed with CO_2_ gas, and the whole eyes (right and left) were enucleated and fixed in a single Eppendorf tube filled with 4% paraformaldehyde (PFA) for 1 h. The eyes were dissected under a stereo zoom microscope (Motic, SMZ-168) to separate the retina from the RPE/choroid. Each retina was washed three times with PBS for 5-10 min. The separated retinal tissues were treated with blocking solution for 1h at room temperature. The blocking solution consisted of 10% normal goat serum (Invitrogen PCN5000), 80% 1X PBS prepared from 10X PBS (Corning, Sigma-Aldrich 46-013-CM), 10% Triton X-100 prepared from Triton X-100 (Sigma T8787).

Separated retinas were then stained with staining solution overnight (8-12 h) at 4 LC. The staining solution consisted of 5% normal goat serum (Invitrogen PCN5000), 84.5% 1X PBS prepared from 10X PBS (Corning, Sigma -Aldrich 46-013-CM), 10% Triton X-100 prepared from Triton X-100 (Sigma T8787) and 0.5% *Griffonia simplicifolia* Isolectin GS-IB4 conjugated with Alexa Fluor 568 (Invitrogen 121412). The retina was divided into four quadrants by four radial cuts that were made from the edge to the center and then mounted on a glass slide with a few drops of Fluoromount-G® mounting medium (Southern Biotech, Birmingham, AL, USA). Both eyes were mounted on a labeled glass slide adjacent to each other. Retinal flat-mount images were captured via a confocal microscope (BX50; Olympus, Tokyo, Japan). The avascular areas, and the total retinal areas were measured manually using Image J (v.2018) imaging software, and the areas of avascular retina VO or INV were quantified with Image pro-plus (Media Cybernetics, Rockville, MD, USA) and expressed as a percentage of total retinal area. ^25, 26^

### Extraction of retinal carotenoids

Retinal carotenoids were extracted from pooled retinas of P17 mouse pups whose mothers had started carotenoid supplementation at P1 (n=4 pairs of retina). The samples were placed into 1 mL tetrahydrofuran (THF) + 0.1% butylated hydroxytoluene (BHT) and 200 μL ethanol and sonicated in an ice-cold water bath for 10 minutes. The extract was then vortexed and centrifuged at 2500 × *g* for 5 min, and the supernatant was collected in a separate tube. The extraction step was repeated two to three more times until the solvent turned colorless. All supernatants were pooled and then evaporated under nitrogen gas at 45°C on a Glas-Col evaporator system (Terre Haute, IN, USA). The dried residue was dissolved in HPLC mobile phase which was then centrifuged at 2500 × *g* for 5 min, and the supernatant was injected into an HPLC system for separation and quantification as previously described. ^23^

### Statistics

Data are presented as mean ± SD. Statistical comparisons were made using a one-way ANOVA followed by Student’s t-tests and Tukey’s multiple comparison tests. Both eyes were imaged, but for scoring and statistical analysis, we typically chose one randomly selected eye per pup unless that eye had an unusable image.

## Results

### Retinal bioavailability of maternally administered carotenoids in Bco2^-/-^ mouse pups

WT mice do not accumulate orally administered L or Z in their retinas even in the setting of high-dose supplementation, but we have previously demonstrated that *Bco2*^*-/-* “^macular pigment mice” take up L and Z into their retinas when they are provided in DSM Actilease® beadlets mixed with their normal diet at a dose of 2.6 mg/day/mouse. ^22^ It has not been shown, however, that maternally administered L or Z can be delivered to their pups’ retinas. The retinas of newborn P1 mouse pups are too small to detect carotenoids levels in the range necessary for HPLC to prove placental carotenoid transmission, so we measured levels from larger P17 mouse retinas to confirm retinal delivery via breastmilk. **Figure 1** examines retinal carotenoid bioavailability in pups nursed by their *Bco2*^*-/-*^ and WT mouse mothers were fed with either placebo, L, or Z supplemented chow starting at P1 and then subjected to our standard OIR protocol. At P17, WT mouse pups never had any detectable retinal carotenoids in any of the supplementation groups. P17 *Bco2*^*-/-*^ mouse pups had no detectable retinal carotenoids when their mothers were fed placebo beadlet chow, while L and Z supplementation resulted in retinal accumulation of 0.103 ng/mouse and 0.125 ng/mouse for L and Z, respectively.

**Figure 1.**
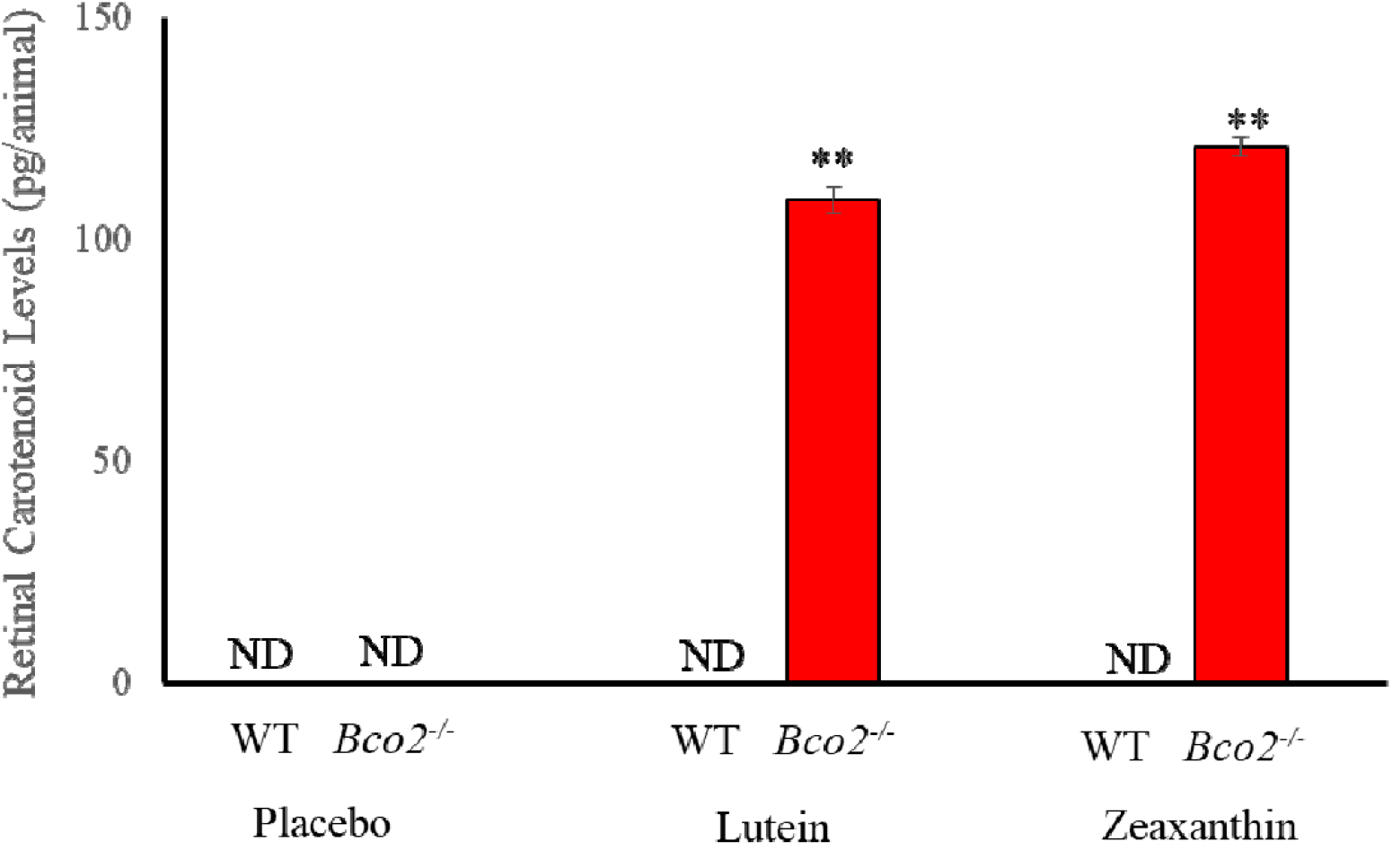
Retinal carotenoids in WT and *Bco2*^*-/-*^ mice subjected to OIR where the nursing mother along with her pups were fed with either lutein, zeaxanthin, or placebo diet starting at postnatal day 1 (P1) and sacrificed on P17. Retinal carotenoids were never detectable in WT mice. Value indicate means ± SD for 3 batches of 8 pooled eyes for each subgroup. ND, not detectable. ** *P*< 0.01 compared to the placebo-fed group and to the WT controls.

### OIR studies in Bco2^-/-^ mouse pups

Nursing mothers along with their litters were subjected to a hyperoxic environment (75% O_2_) from P7-12 in the OIR chamber causing VO in the central retina of the pups. The central avascular retinal area was measured at P12. **Figure 2** shows that pups of *Bco2*^*-/-*^ mothers fed prenatally with a placebo diet starting at E0 (equivalent to the beginning of the first trimester of human pregnancy) exhibit a significantly higher percentage of VO (28.3%) compared to the L-fed group (10.7 %) and the Z-fed group (8.6 %). Likewise, pups of *Bco2*^*-/-*^ mothers fed with the placebo diet starting at E11 (equivalent to the beginning of the second trimester of human pregnancy) have a significantly higher percentage of VO (27.7%) relative to the L-fed group (12.6%) and the Z fed group (10.3%) (**Figure 3**). When maternal carotenoid supplementation was initiated postnatally at P1, a timepoint in mouse retinal development equivalent to the beginning of the third trimester of human pregnancy, L and Z were still significantly protective against OIR VO (26.2%, 20.2%, and 18.9% for placebo, L, and Z, respectively), but with less efficacy (**Figure 4**). A replot of the data from these experiments in **Figure 5** demonstrates that prenatally initiated supplementation with L or Z was always significantly better than postnatally initiated supplementation. There was a consistent trend that Z supplementation was superior to L supplementation, but this was not statistically significant.

**Figure 2.**
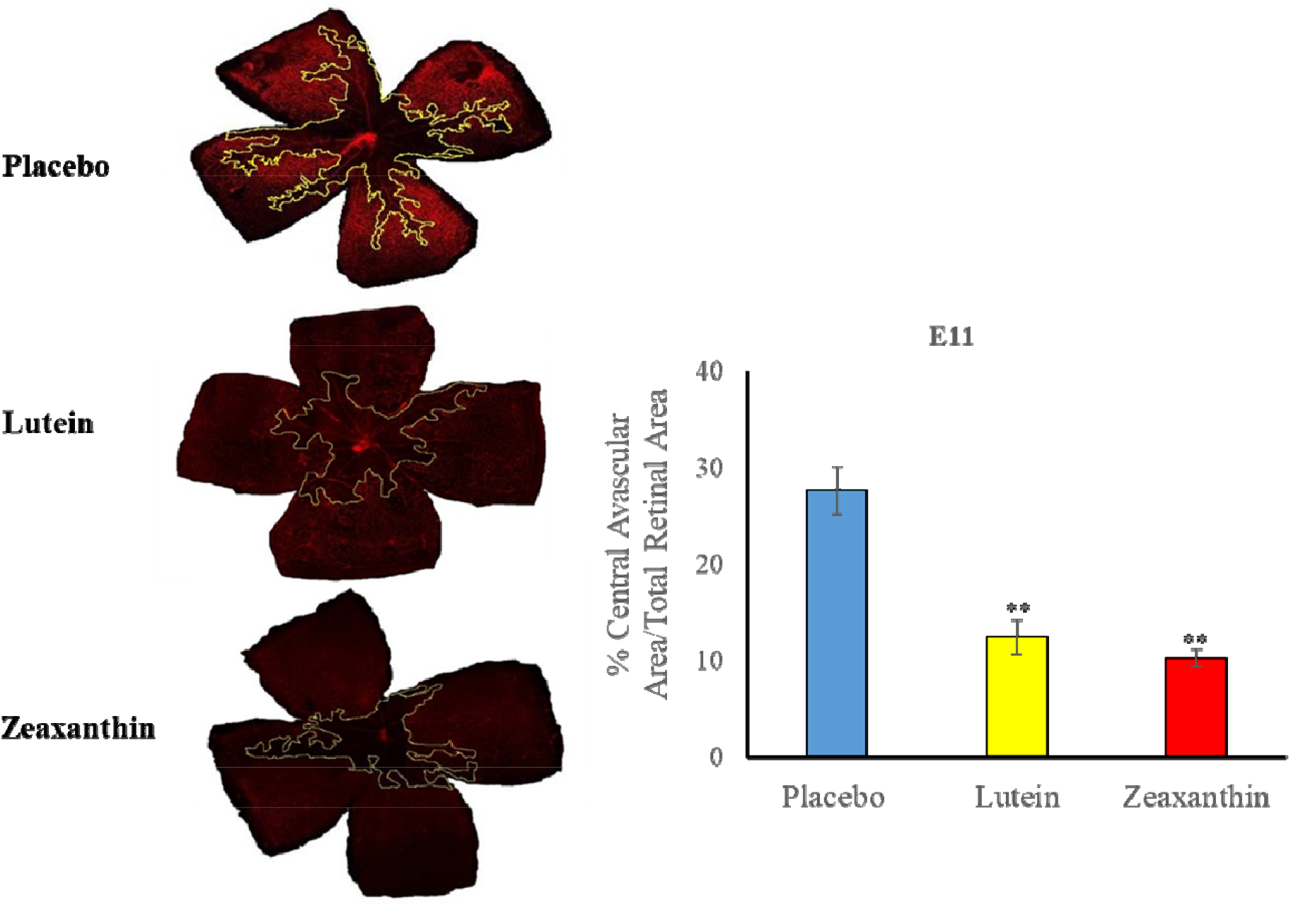
Representative lectin-stained flatmounts of retinas from *Bco2*^*-/-*^ mouse pups sacrificed at P12 showing vaso-obliteration (VO) after treatment under our OIR protocol. Their mothers started supplementation with L, Z, or placebo diets at E0. The yellow marking shows the central avascular area. ** *P*< 0.01, compared to the placebo group. Placebo: 6 pups from 2 litters; Lutein: 4 pups from 4 litters; Zeaxanthin: 4 pups from 3 litters.

**Figure 3.**
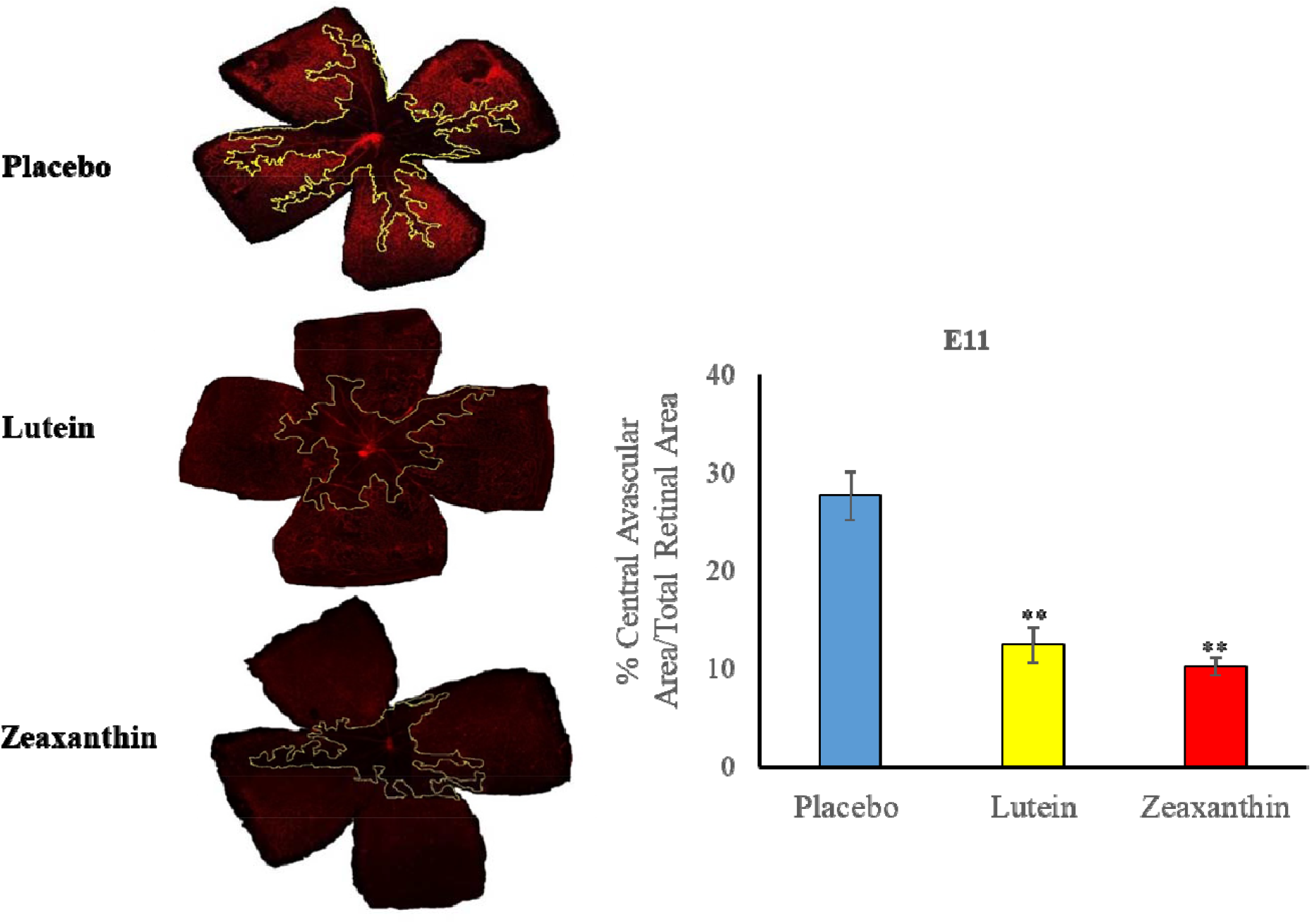
Representative lectin-stained flatmounts of retinas from *Bco2*^*-/-*^ mouse pups sacrificed at P12 showing VO after treatment under our OIR protocol. Their mothers started supplementation with L, Z, or placebo diets at E11. The yellow marking shows the central avascular area. ** *P*< 0.01, compared to the placebo group. Placebo: 5 pups from 2 litters; Lutein: 5 pups from 3 litters; Zeaxanthin: 5 pups from 3 litters.

**Figure 4.**
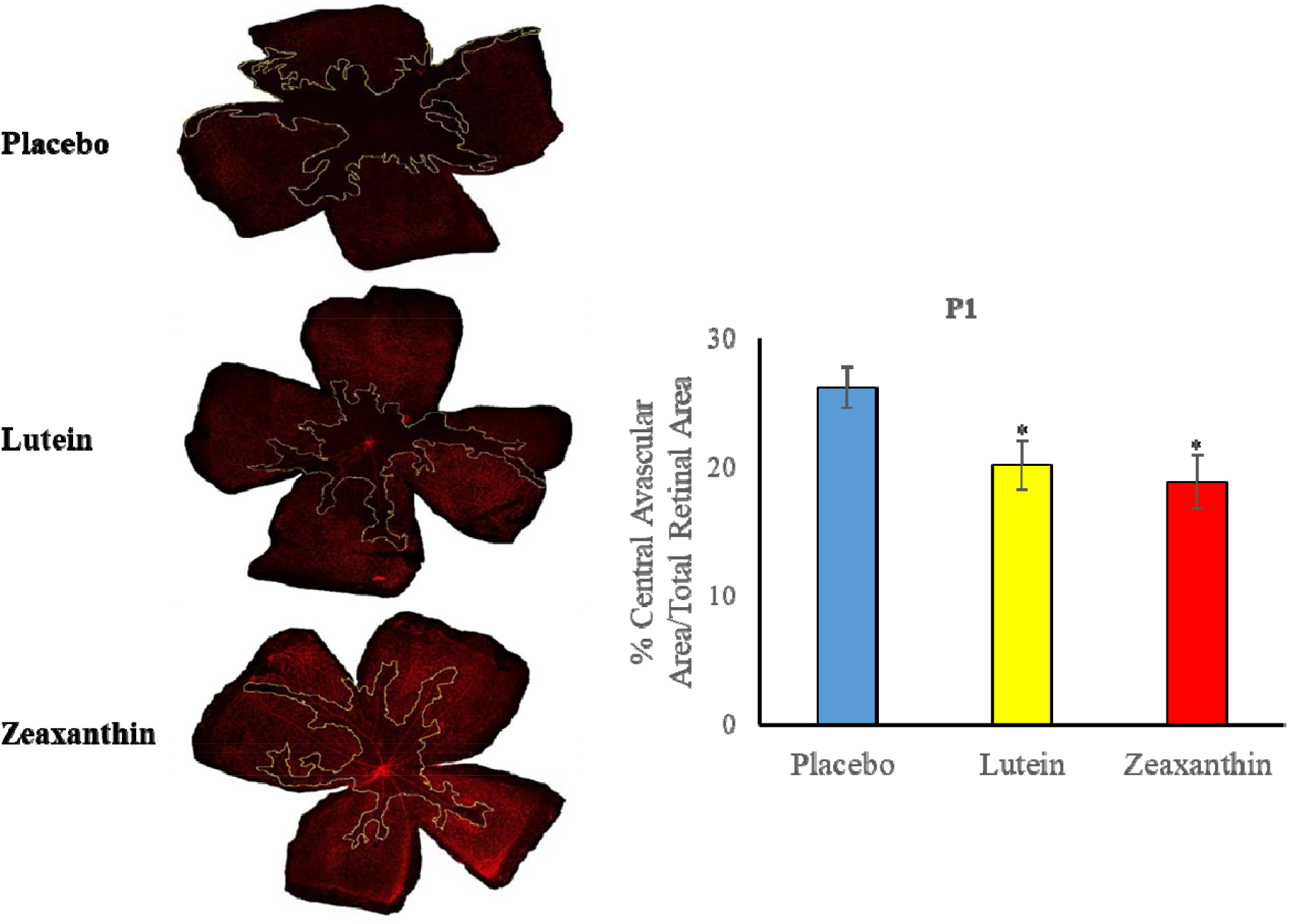
Representative lectin-stained flatmounts of retinas from *Bco2*^*-/-*^ mouse pups sacrificed at P12 showing VO after treatment under our OIR protocol. Their mothers started supplementation with L, Z, or placebo diets at P1. The yellow marking shows the central avascular area. * *P*< 0.05, compared to the placebo group. Placebo: 6 pups from 2 litters; Lutein: 5 pups from 2 litters; Zeaxanthin: 5 pups from 2 litters.

**Figure 5.**
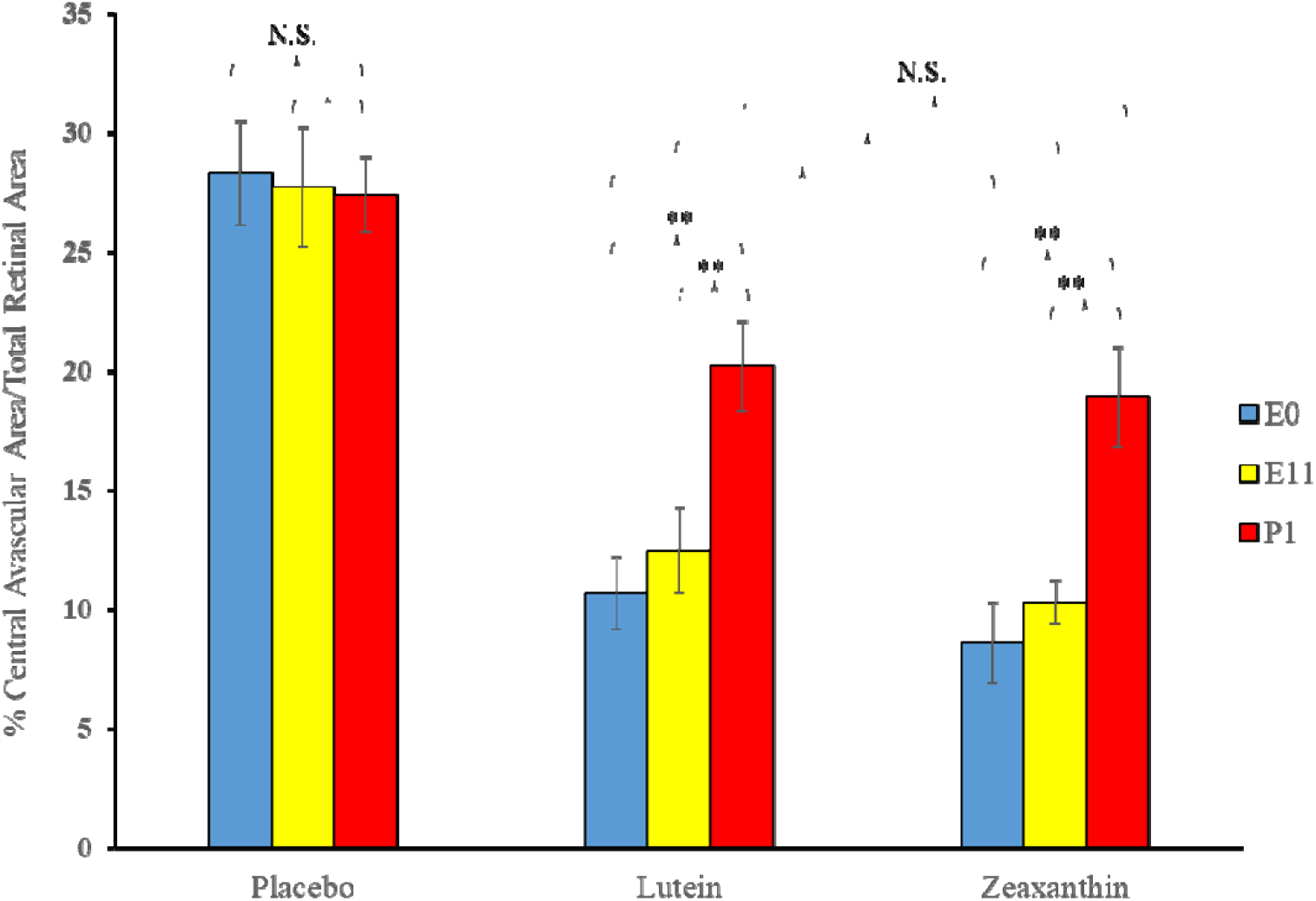
Replot of the VO results from the OIR experiments of **Figures 2-4**. Pregnant *Bco2*^*-/-*^ mice started L, Z, or placebo supplementation on E0, E11, or P1, time points in mouse gestation corresponding to the three trimesters of retinal development during human pregnancy. ** *P*< 0.01, compared to the postnatal group, P1; NS: not significant.

When nursing *Bco2*^*-/-*^ mothers along with their litters were moved from the hyperoxia environment (75% O_2_) on P12 to room air (21% O_2_) for 5 days, neovascular tufts formed corresponding to the second phase of human ROP. **Figures 6-8** show the INV area of retinal flat mounts of P17 *Bco2*^*-/-*^ mouse pups whose mothers were fed with either placebo, L, or Z diets starting at E0, E11, or P1. Compared to the placebo-fed groups that demonstrated 8.3% INV, groups fed prenatal supplementation of L and Z starting at E0 had significantly reduced retinal INV at 4.5% and 3.0%, respectively **(Figure 6)**. Likewise, prenatal supplementation of L and Z at E11 to pregnant mothers had significantly reduced retinal INV area at 5.0% and 4.1%, respectively, compared to the placebo-fed groups at 7.9% **(Figure 7)**. Pups of *Bco2*^*-/-*^ mouse mothers postnatally fed with placebo diet starting at P1 had a significantly higher percentage of INV (7.6%) compared to the L fed group (5.7%) and the Z fed group (5%), respectively **(Figure 8)**. Similar to our VO experiments, a replot of the data from these INV experiments in **Figure 9** demonstrates that prenatally initiated supplementation with L or Z was always better than postnatally initiated supplementation, but the improvement was statistically significant only if carotenoid supplementation was initiated at E0. Again, there was a consistent trend that Z supplementation was superior to L supplementation, but this was not statistically significant.

**Figure 6.**
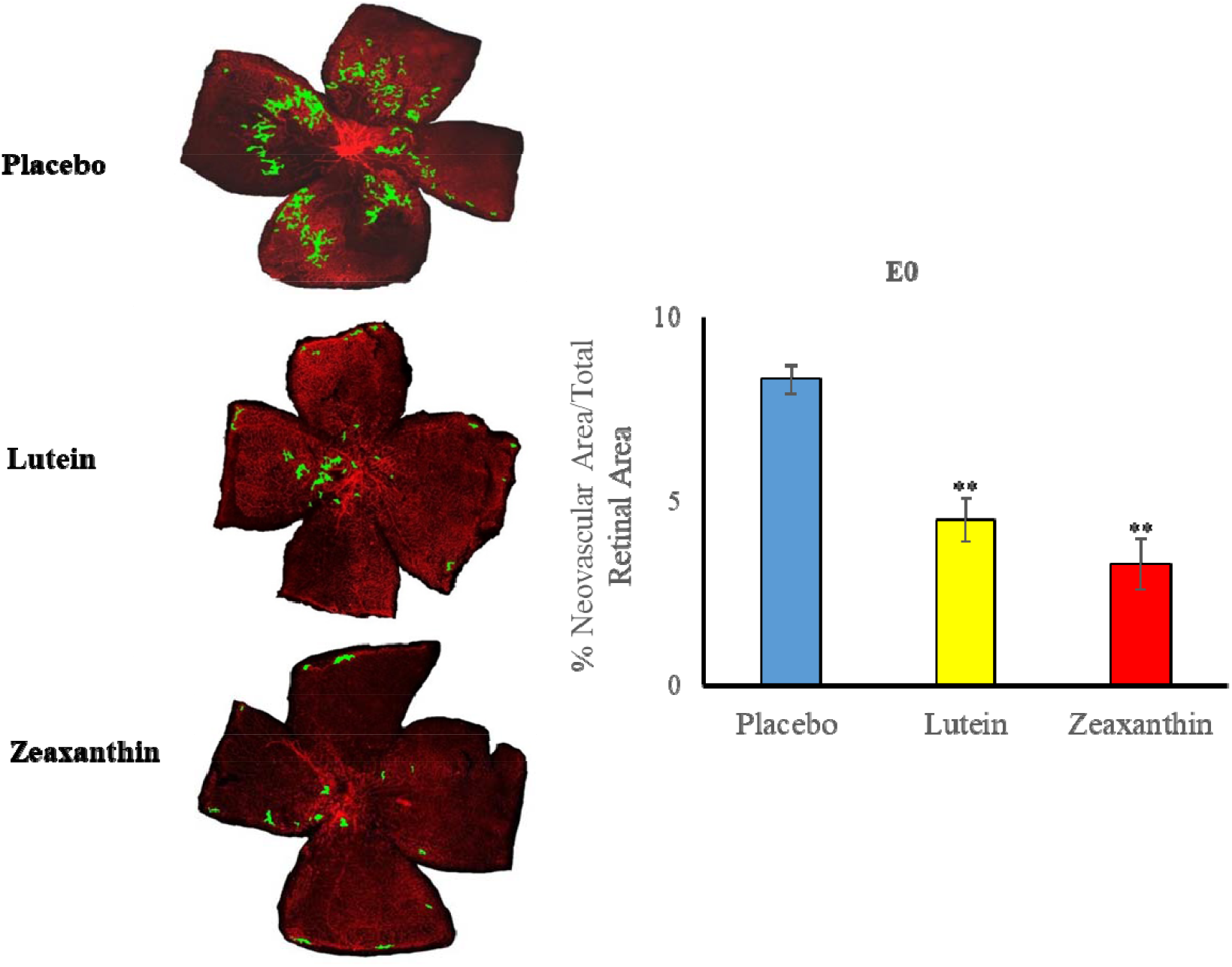
Representative lectin-stained flatmounts of retinas from *Bco2*^*-/-*^ mouse pups sacrificed at P17 showing intravitreal neovascularization (INV) after treatment under our OIR protocol. Their mothers started supplementation with L, Z, or placebo diets at E0. The green marking shows the INV area. ** *P*< 0.01, compared to the placebo group. Placebo: 5 pups from 3 litters; Lutein: 3 pups from 3 litters; Zeaxanthin: 3 pups from 3 litters.

**Figure 7.**
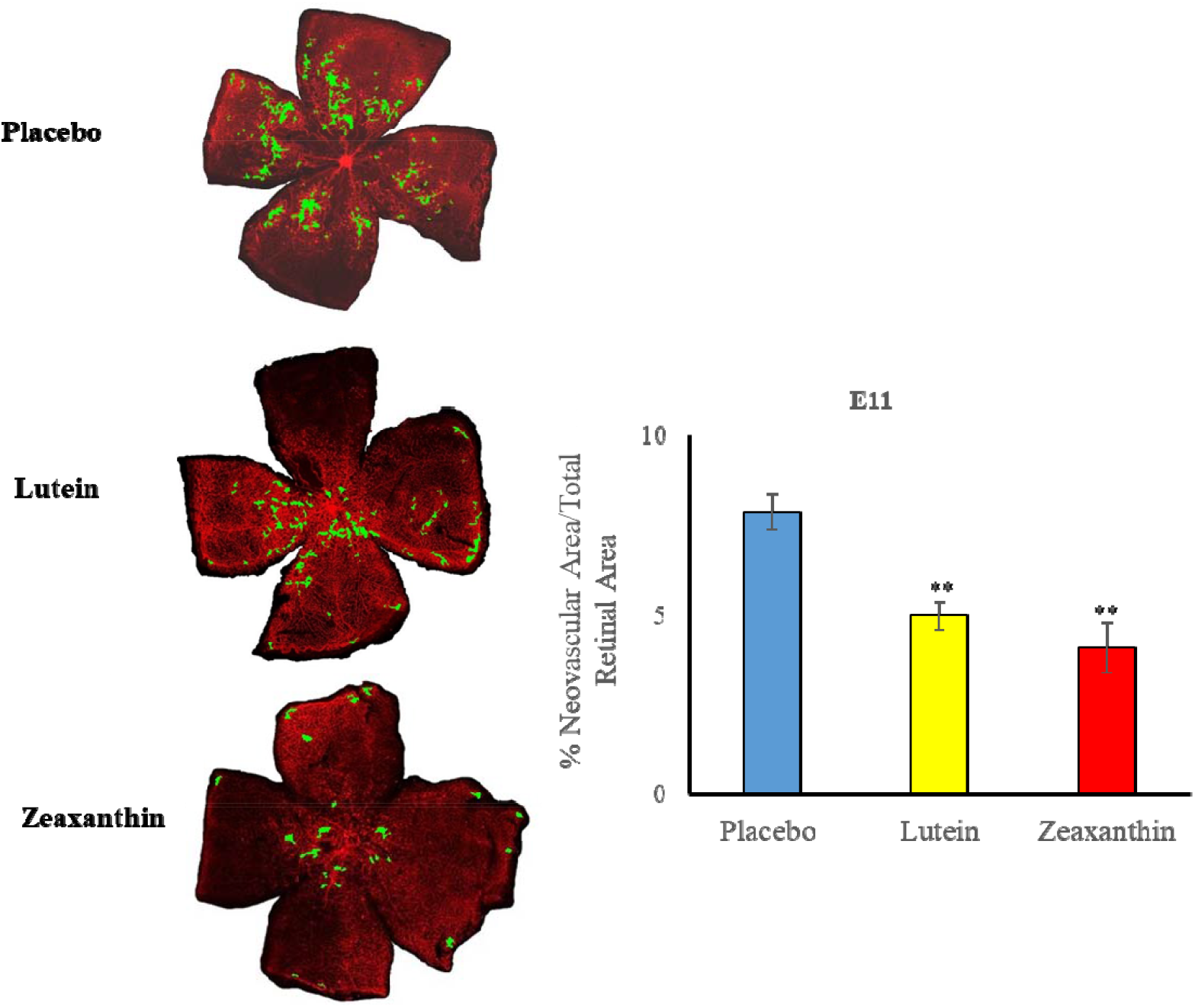
Representative lectin-stained flatmounts of retinas from *Bco2*^*-/-*^ mouse pups sacrificed at P17 showing INV after treatment under our OIR protocol. Their mothers started supplementation with L, Z, or placebo diets at E11. The green marking shows the INV area. ** *P*< 0.01, compared to the placebo group. Placebo: 5 pups from 2 litters; Lutein: 4 pups from 2 litters; Zeaxanthin: 4 pups from 2 litters.

**Figure 8.**
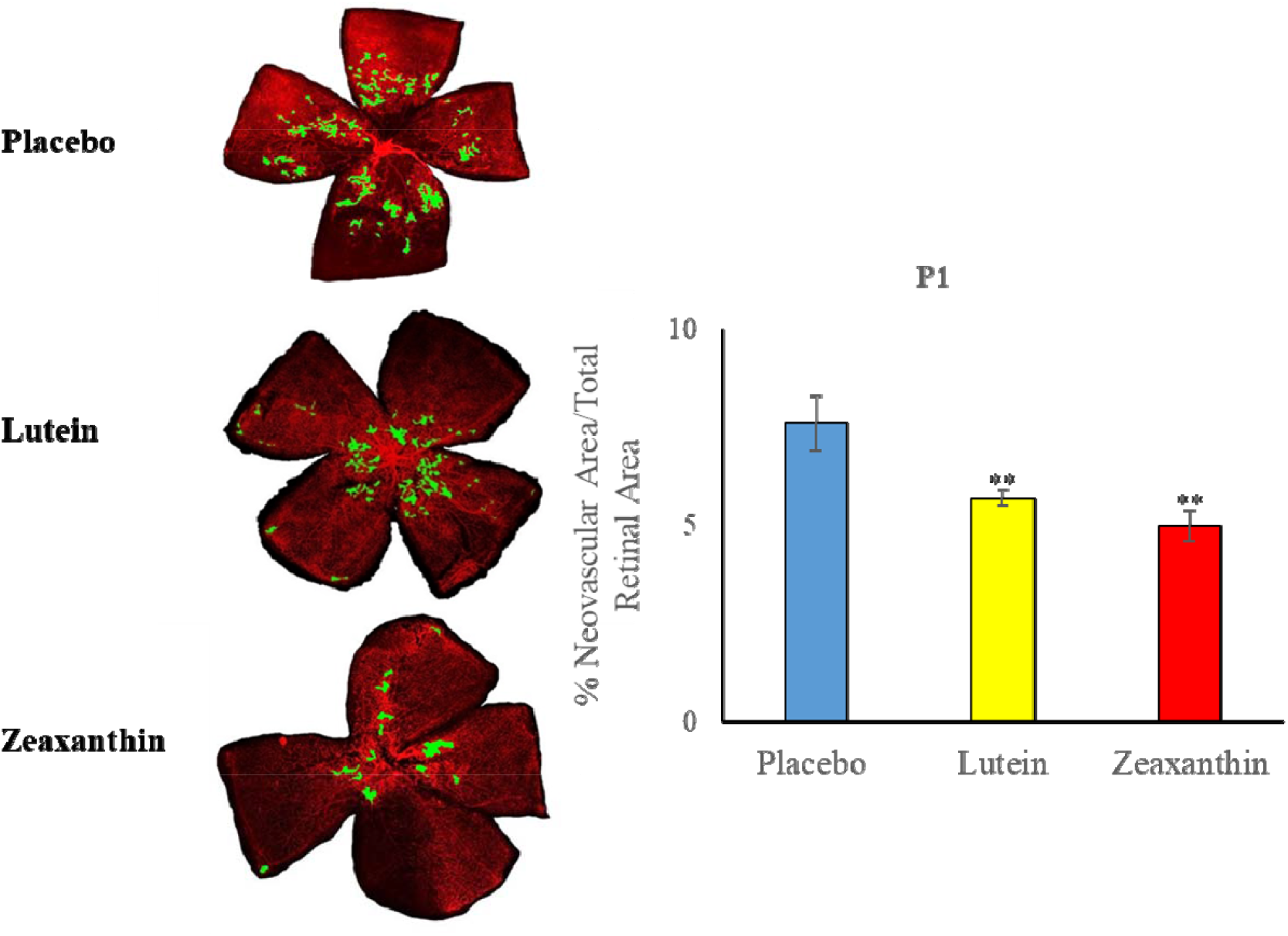
Representative lectin-stained flatmounts of retinas from *Bco2*^*-/-*^ mouse pups sacrificed at P17 showing INV after treatment under our OIR protocol. Their mothers started supplementation with L, Z, or placebo diets at P1. The green marking shows the INV area. ** *P*< 0.05, compared to the placebo group. Placebo: 5 pups from 2 litters; Lutein: 5 pups from 2 litters; Zeaxanthin: 5 pups from 2 litters.

**Figure 9.**
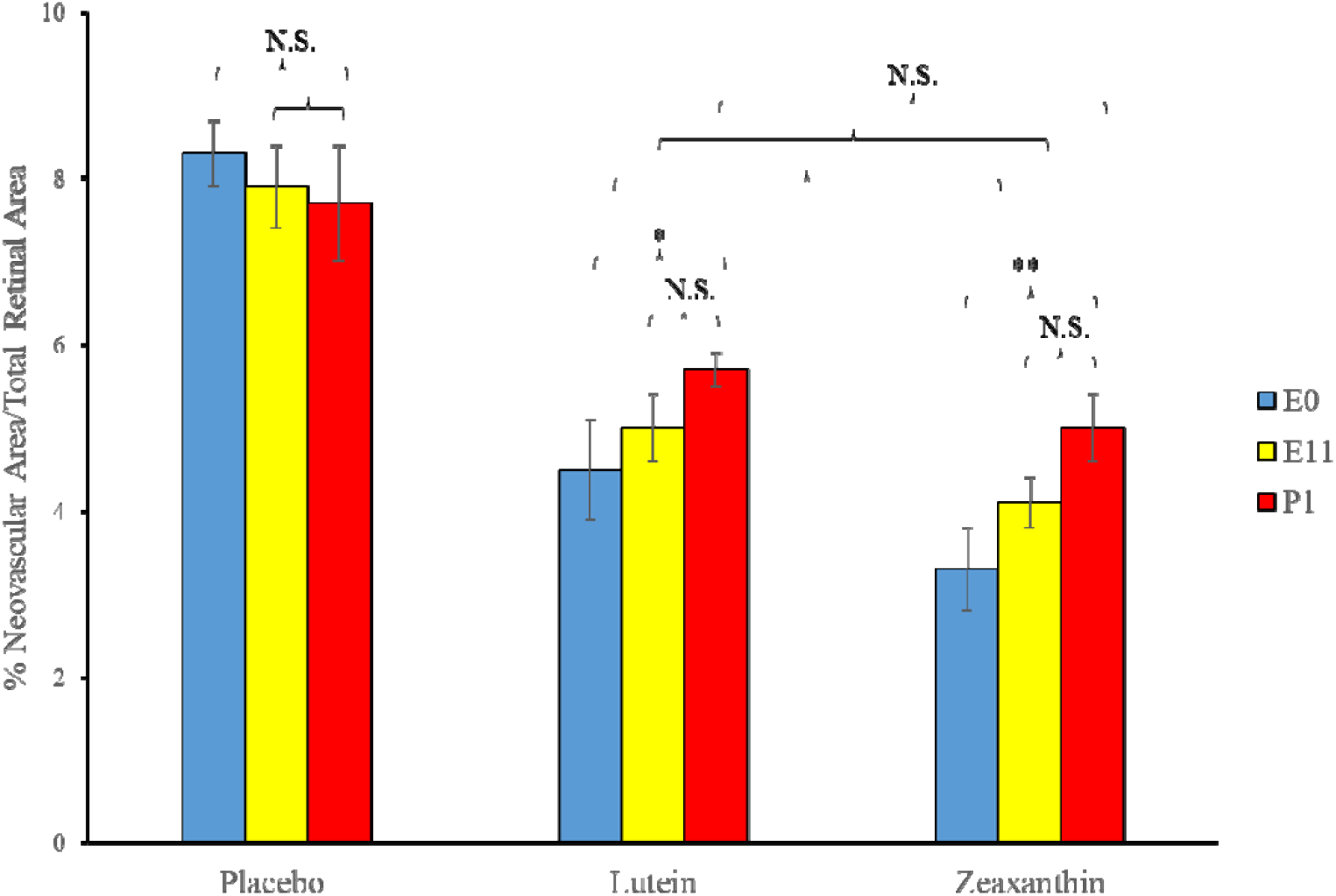
Replot of the INV results from the OIR experiments of **Figures 6-8**. Pregnant *Bco2*^*-/-*^ mice started L, Z, or placebo supplementation on E0, E11, or P1, time points in mouse gestation corresponding to the three trimesters of retinal development during human pregnancy. * *P*< 0.05, ** *P*< 0.01, compared to the P1 postnatal group; NS: not significant.

## Discussion

In designing this study, we hypothesized that maternal prenatal supplementation with eye-protective carotenoids may help to prevent or ameliorate human ROP and tested this hypothesis in a mouse model of OIR. This hypothesis was based on the macular carotenoids’ specific prenatal deposition in the developing human retina, their well-known antioxidant properties, and the high levels of oxidative stress in premature infants who have not benefited from the third trimester’s placental transfer of L and Z from the mother to her child. Recent recognition that WT mice are a suboptimal model for diet-based carotenoid interventions against ocular disease guided us to use *Bco2*^*-/-*^ mice that reproducibly deposit L and Z in their retinas when supplemented with these MP carotenoids. ^22^

Oxidative stress is thought to be a major etiological factor for ROP. The newborn ICU is a stressful environment where high levels of oxygen, light, and high-intensity medical and surgical interventions are routinely implemented on infants with extremely low systemic and ocular carotenoid levels and immature antioxidant-based defense systems. ^27^ Thus, it is not surprising that the oxidative stress marker 8-hydroxy-2-deoxyguanosine (8-OHdG) has been reported to be at a higher concentration in leukocytes and urine of ROP infants compared to full-term infants. ^28^ Prenatal carotenoid supplementation in newborn infants has been reported to decrease plasma oxidative stress biomarkers and increase antioxidant enzyme levels. ^27,29^ Excessive oxygen or fluctuations in oxygen levels are believed to be involved in the first phase of ROP where there is delayed normal retinal vascular growth and damage to existing capillaries. When premature babies are transferred to room air after oxygen therapy, the avascular retina becomes hypoxic, releases even more reactive oxygen species (ROS), and upregulates angiogenic factors leading to pathological intravitreal neovascularization characterized by neovascular tufts at the junction of the avascular and vascular zones. ^30^

Minimizing ROS and oxidative stress-induced damage in preterm infants through maternal prenatal supplementation with carotenoids could be an effective nutritional therapy against ROP. The macular antioxidant carotenoids, L and Z, protect the retina from oxidative stress and light-induced damage and can enhance visual performance in adults. ^31, 32^ These eye-protective nutrients begin to accumulate in the developing retina during the second trimester of pregnancy and are actively transported in such large amounts from mother to infant during the third trimester that the mother’s systemic carotenoid levels may even decline. We have shown both cross-sectionally and in the recently completed L-ZIP randomized clinical trial that the mother’s ocular and systemic carotenoid status strongly influences her newborn’s ocular and systemic carotenoid status. ^18^ Moreover, L and Z supplementation at AREDS2 dosages was safe and well-tolerated during pregnancy.

The present study demonstrates that *Bco2*^*-/-*^ mouse pups whose mothers received oral MP carotenoid supplementation accumulated L and Z in their retinas, while WT pups did not. Both prenatal and postnatal supplementation significantly inhibited hyperoxia-induced VO and hypoxia-induced INV OIR in *Bco2*^*-/-*^ pups, but prenatal supplementation was more effective than postnatal initiation of supplementation in this mouse model of OIR. Both macular carotenoids attenuated OIR pathology but did not affect normal retinal vascular growth or development. L and Z were effective in protecting against OIR pathology not only in the hyperoxia phase (P7-P12) but also extended protection into the hypoxia phase (P12-P17). Z was somewhat more effective than L in retinal protection against OIR complications in our *Bco2*^*-/-*^ mice, possibly because Z has better bioavailability relative to L in mice ^23^ and because Z has higher antioxidant activity than L due to its longer conjugated double-bond structure. ^33^ Similarly, dietary supplementation with Z in Japanese quail exhibited a better photo-protective effect than L against light-induced photoreceptor damage. ^34^

The molecular mechanisms underlying the protective effect of carotenoids against OIR still need to be studied in detail. L and Z may protect against and ameliorate OIR by controlling vessel loss in an oxygen-stressed environment by reducing the hypoxic stimulus for pathological INV. Studies on the *apoE*^*-/-*^ mouse model for AMD-like retinal degeneration and on the streptozotocin-induced diabetic retinopathy mouse model have shown that L treatment downregulates VEGF expression and elevates antioxidant enzymes levels. ^35, 36^

Previously published studies of the potential benefits of L and Z for the prevention and treatment of ROP and OIR have had disappointing results. Even though an AREDS2-type dosage of ∼0.14 mg/kg/day was used in human premature infant studies, supplementation was started only after birth, and systemic bioavailability of the formulation was not confirmed. ^19, 20, 21^ Fu *et al*. (2017) used WT mice supplemented daily by intraperitoneal administration of 0.2 mg/kg of L in 10% DMSO from P7 to P11 in a mouse model of OIR and found no significant changes in retinal VO or INV area relative to control administration of 10% DMSO. ^37^ Our contrasting positive results presumably arise from starting our intervention prenatally and the use of *Bco2*^*-/-*^ mice that are known to take up carotenoids into their retinas because their highly active xanthophyll cleavage enzyme has been knocked out.

The results of this animal study, coupled with our L-ZIP clinical study results showing the safety and biological efficacy of carotenoid supplementation during pregnancy, suggest that the potential benefits of L and Z against ROP should be revisited, this time in a prospective, randomized clinical trial of prenatal maternal supplementation as opposed to the prior human and animal studies which have focused on postnatal supplementation. Even though there are currently no prenatal vitamins or postnatal premature infant formulas on the market with added L and Z, they can be readily prepared and appear to be safe and well tolerated. ROP continues to be a major public health problem worldwide, and a proven, low risk, relatively inexpensive nutritional intervention would be welcome.

## Acknowledgments

We would like to thank Dr. Johannes von Lintig from Case Western Reserve University for generously providing the *Bco2*^*-/-*^ founder mice.

---

**Supplementary Table 1.** Schematic representation of animal experimental design showing different treatment groups with their respective timelines and endpoints.

| **Timeline** | **Pregnant *Bco2*<sup>-/-</sup> mice (n = 2-4 mothers in each experimental group)** |  |  |
| --- | --- | --- | --- |
| **Prenatal L, Z, and placebo supplementation start at different pregnancy stages.** | **Group I**<br>**E0 – Embryonic day 0**<br>**(~ 1<sup>st</sup> trimester in humans)** | **Group II**<br>**E11 – Embryonic day 11**<br>**(~ 2<sup>nd</sup> trimester in humans)** | **Group III**<br>**P1 – Postnatal day 1**<br>**(~ preterm baby in humans)** |
|  | **Each group was subdivided into the L, Z, and placebo subgroups**<br>• Pups from each experimental subgroup were pooled from 2-4 mothers.<br>• Pups' body weights were assessed at P7, P12, and P17. |  |  |
| **P7 – P12**<br>**Hyperoxia phase** | • Nursing mothers and pups were exposed to 75% O<sub>2</sub> to initiate OIR<br>• At the end of this phase, pups (n=4-6/subgroup) were sacrificed, and the retinas were analyzed for vaso-obliteration (VO). |  |  |
| **P12 –17**<br>**Hypoxia phase** | • Nursing mothers and the remaining pups were returned to 21% O<sub>2</sub>.<br>• At the end of this phase, the pups (n=3-5/subgroup) were sacrificed, and the retinas were analyzed for intravitreal neovascularization (INV). |  |  |

**Supplementary Figure 1.**
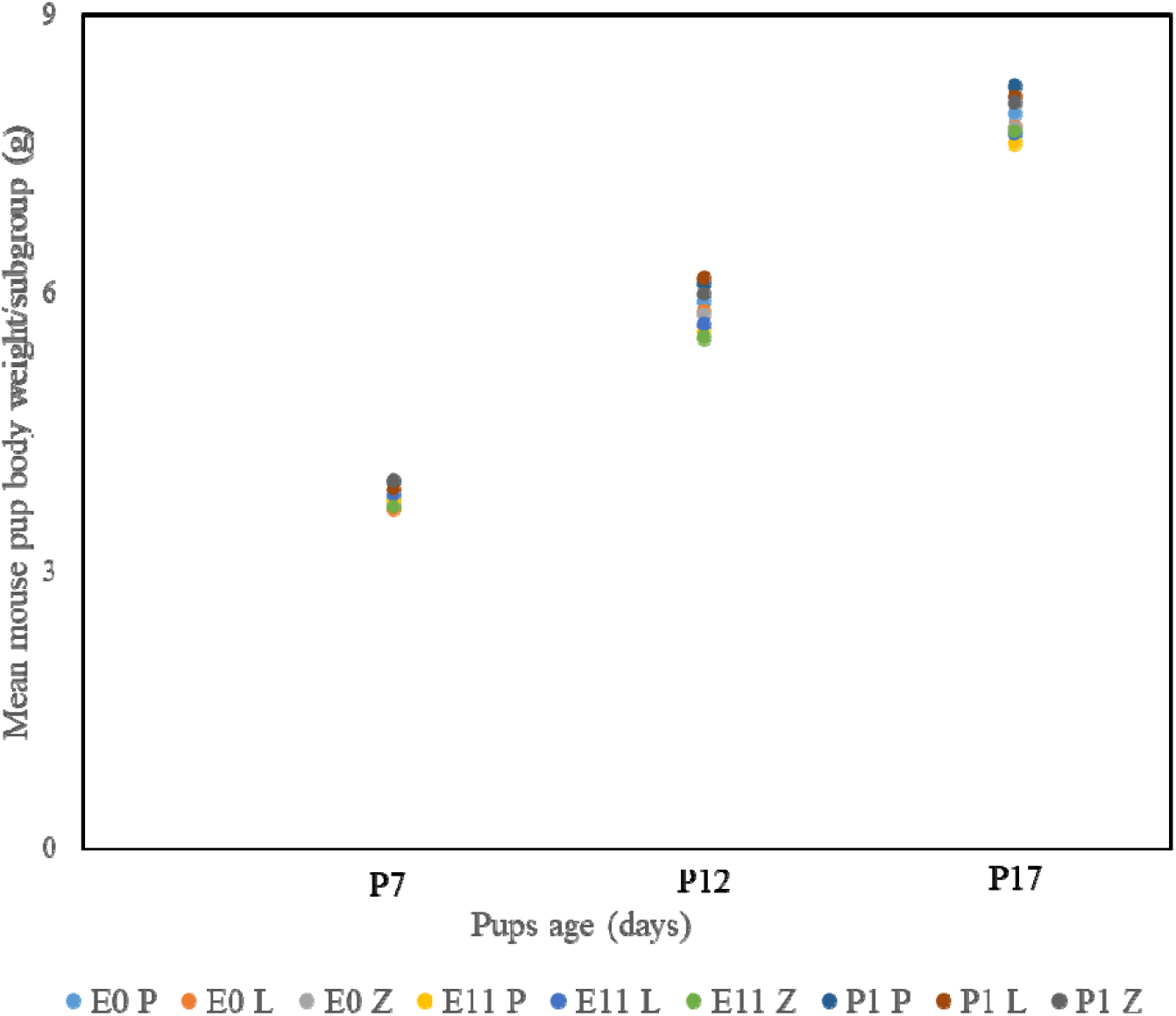
Mean body weight gain of *Bco2*^*-/-*^ mouse pups whose pregnant mothers were supplemented with either lutein (L), zeaxanthin (Z), or placebo (P) diets starting at E0, E11, or P1 and then subjected to our OIR protocol. Their body weights were measured at P7, P12, and P17.

## Notes

**Conflict of Interest:** No authors have any conflicts of interest.

**Grant Support:** NEI grants EY11600 (PSB), EY14800 (PSB), EY15130 (MEH), EY17011 (MEH), and an unrestricted grant from Research to Prevent Blindness to the Department of Ophthalmology and Visual Sciences.

### Competing Interest Statement

The authors have declared no competing interest.

